# N’-(1-phenylethylidene)-benzohydrazide cytotoxicity is LSD1 independent and linked to Fe-S cluster disruption in Ewing sarcoma

**DOI:** 10.1101/2025.06.20.660795

**Authors:** John W. Sherman, Galen C. Rask, Bingcong Xiong, E. John Tokarsky, Aundrietta Duncan, Eranthie Weerapana, Emily R. Theisen

**Author notes:** To whom correspondence should be addressed. Abigail Wexner Research Institute, Nationwide Children’s Hospital, 575 Children’s Crossroads, Columbus, OH, 43215. **Disclosures:** E.R.T. has received research funding in the past from Salarius Pharmaceuticals that is related to the work performed here. E.R.T. currently receives no research funding from Salarius Pharmaceuticals and has no financial stake in the company. A.D. declares competing interests as an employee of Salarius Pharmaceuticals during the time the project was being performed. No disclosures are reported by other authors.

## Abstract

The noncompetitive LSD1 inhibitors SP-2509 and SP-2577 are N’-(1-phenylethylidene)-benzohydrazides that display potent activity in Ewing sarcoma. They block transcriptional regulation of the causative oncogenic fusion protein, EWSR1::FLI1, and cause cell death. However, SP-2509 and SP-2577 are the only LSD1 inhibitors active in Ewing sarcoma; other LSD1 inhibitors have little effect. Studies from our group and others suggest SP-2509 activity may result from off-target activity affecting the mitochondria. Here we identified potential off-target mechanisms of N’-(1-phenylethylidene)-benzohydrazides using an unbiased approach, cellular thermal shift assay coupled to mass spectrometry (CETSA-MS). Interestingly, this revealed significant destabilization of the electron transport chain complex III protein ubiquinol-cytochrome c reductase (UQCRFS1). We find that UQCRFS1 destabilization is likely linked to impaired iron-sulfur (Fe-S) cofactor binding, and that SP-2509 broadly destabilizes cellular Fe-S proteins. Using both chemical and genetic tools, we show that SP-2509 mediated cell death is LSD1 independent and instead requires a N’-(2-hydroxybenzylidene)hydrazide. Our studies suggest this core moiety alters iron metabolism in the cell. Importantly, we also find that the reversal of EWSR1::FLI1 transcriptional regulation by SP-2509 is independent from LSD1 inhibition. This unique activity is instead associated with the N’-(2-hydroxybenzylidene)-hydrazide core and destabilization of Fe-S proteins. These findings reveal a novel mechanism of action for this class of compounds and raise additional questions regarding how EWSR1::FLI1 transcriptional regulation is linked to Fe-S biogenesis, the precise mechanisms of cell death, the biological features of susceptible cancer cells, and strategies for clinical translation.

## INTRODUCTION

The molecules SP-2509 and SP-2577 (seclidemstat) are N’-(1-phenylethylidene)-benzohydrazides that were first described as inhibitors of the lysine specific demethylase 1 (LSD1) in 2013 (Fig. 1A). ^1^ This class of molecules has shown promise for the treatment of several cancers, with seclidemstat entering clinical trials for Ewing sarcoma and related *FUS*, *EWSR1*, and *TAF15* (FET)-rearranged tumors (NCT03600649, NCT05266196), as well as myelodysplastic syndrome (MDS) and chronic myelomonocytic leukemia (CMML) (NCT04734990). In Ewing sarcoma, treatment with SP-2509 and SP-2577 blocks oncogenic transcriptional regulation mediated by the causative fusion protein, EWSR1::FLI1, and leads to cell death.^2^ LSD1 colocalizes with EWSR1::FLI1 throughout the genome, and overexpression in Ewing sarcoma is associated with poor prognosis in patients. Both shRNA-mediated LSD1 knockdown and EWSR1::FLI1 knockdown cause similar changes in the transcriptome as measured by RNA-seq. ^2–4^

**Figure 1.**
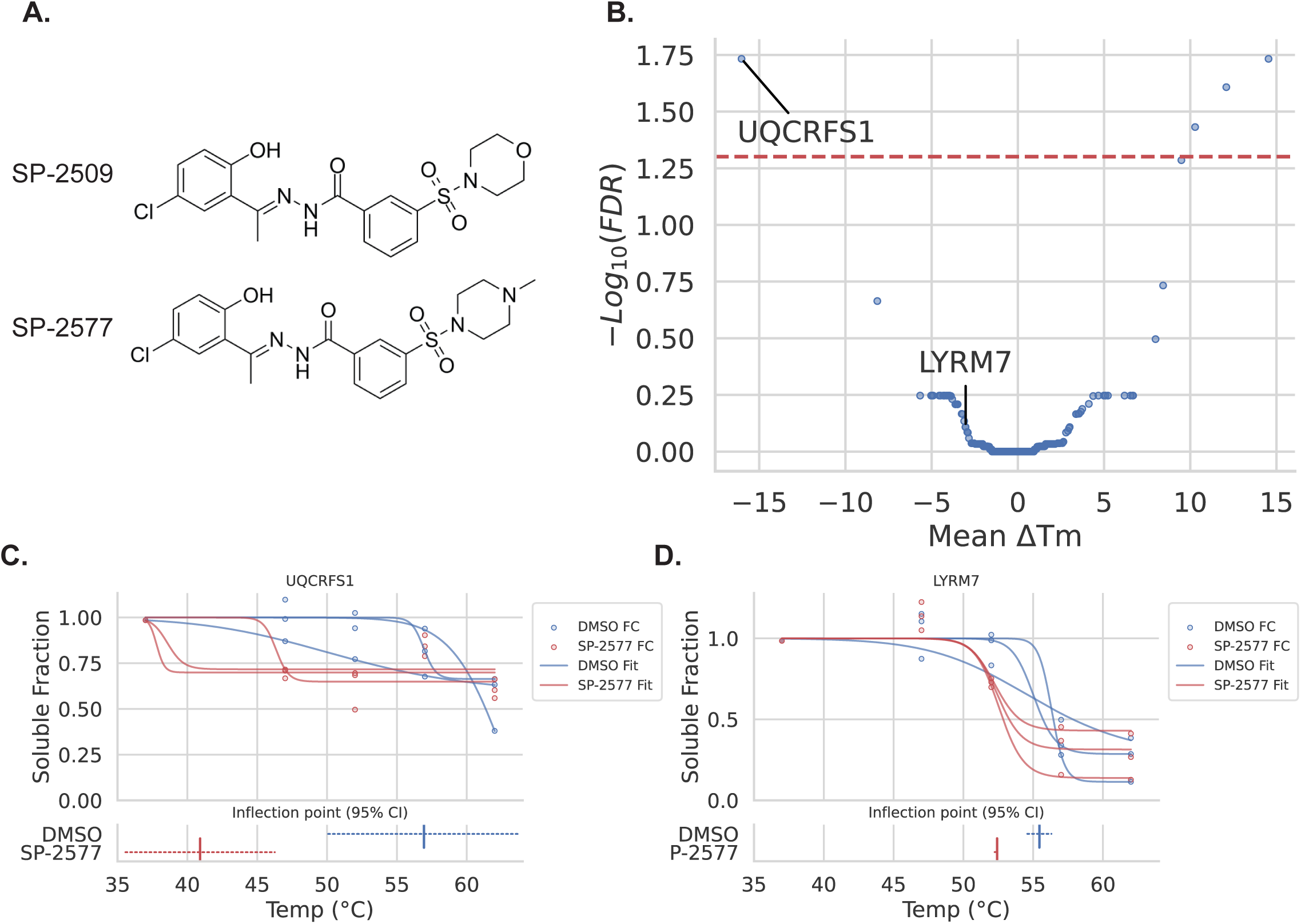
CETSA-MS identifies UQCRFS1 as destabilized by N’-(2-hydroxybenzylidene)hydrazide treatment. A) Chemical structure of SP-2509 and SP-2577. TMT-labeled CETSA mass spectrometry was performed on A673 cells treated with either 0.1% DMSO or 1 µM SP-2577 for 3 hrs. B) Volcano plot of mean ΔTm (Tm SP-2577 – Tm DMSO, N = 3) vs -Log10(FDR) (bootstrap with Benjamini-Hochberg correction). Red line indicates FDR < 0.05 cuttoff. C, D) Graphs of modeled melt curves for UQCRFS1 and LYRM7 protein from. In the top graph, dots represent sample corrected fold change from 37 °C, while lines represent curve of best fit using a 4 parameter logorithmic model. In the lower graph, vertical hashs represent the calculated melting temperature (mean inflection point, N = 3), while dashed line represents 95% confidence interval.

For many years LSD1 inhibition has been considered a promising therapeutic strategy for Ewing sarcoma, however, there are significant differences in the efficacy of the N’-(1-phenylethylidene)-benzohydrazides (SP-2509 and SP-2577) and other LSD1 inhibitors that remain unclear. ^5–8^ SP-2509 is a reversible, noncompetitive LSD1 inhibitor, suggesting an undefined allosteric binding pocket. Other LSD1 inhibitors (including tool compounds and clinical candidates) are predominantly irreversible inhibitors that bind the FAD cofactor or reversible competitive inhibitors that block substrate binding. ^1, 7–9^ Other groups have shown that treatment with SP-2509 and SP-2577 impairs critical protein-protein interactions (PPIs), such that SP-2509 and SP-2577 have been called scaffolding inhibitors. ^10^ The inhibition of LSD1 scaffolding function in combination with a cell state that depends on specific LSD1 PPIs has been proposed as the mechanistic basis for the unique efficacy of noncompetitive LSD1 inhibitors in Ewing sarcoma.

The N’-(2-hydroxybenzylidene)hydrazide core of SP-2509 and SP-2577 is a known pan-assay interference moiety, which may lead to false positive hits in biological assays through a variety of off-target mechanisms. ^11, 12^ Two recent reports using orthogonal approaches to determine cellular mechanisms of resistance to SP-2509 found altered mitochondrial function as a critical feature of drug-resistant cells.^13, 14^ Here, we sought to further define possible N’-(1-phenylethylidene)-benzohydrazide off-target mechanisms that lead to Ewing sarcoma cell death and found an association between cytotoxicity and disruption of proteins requiring Fe-S clusters rather than through inhibition of LSD1 activity.^8, 15^ Using genomics approaches and additional chemical tools we have identified a major mechanism of N’-(1-phenylethylidene)benzohydrazide driven cell death in Ewing sarcoma that is LSD1 independent.

## RESULTS

### SP-2577/2509 alters the stability of several mitochondrial Fe-S cluster proteins

To uncover possible off-target mechanisms of N’-(1-phenylethylidene)-benzohydrazides in Ewing sarcoma cells, we used an unbiased proteomics approach. Shotgun proteomics using mass spectrometry was coupled to the Cellular Thermal Shift Assay (CETSA-MS) to determine candidate N’-(1-phenylethylidene)- benzohydrazide binding partners.^16^ The thermal stabilities of cellular proteins were determined by assaying the melting temperature (Tm) across the proteome after 3 hrs of treatment with either 1 µM SP-2577 or DMSO. A canonical drug target interaction shows that binding of a small molecule results in greater protein stability thus requiring increased temperature to cause unfolding (positive ΔTm value). Conversely, proteins that melt at lower temperatures (negative ΔTm) are destabilized, and this can occur through loss of stabilizing interactions with other proteins, ligands, or cofactors. The majority of cellular proteins experience minimal changes in the calculated Tm (Fig. 1B, Supplemental Table 1) indicating that off-target activity is not broadly observed in the proteome. Notably, our assay identified 5 proteins with increased Tm, and 3 proteins with decreased Tm, using a cutoff of Benjamini–Hochberg FDR < 0.05, indicating SP-2577 off-target activity for these proteins (Fig. 1B).

The second most destabilized protein (decreased ΔTm) was UQCRFS1 (ΔTm = -16.0°C, Fig. 1C), knockout of which was previously reported to cause resistance to SP-2509 in A673 cells. ^14^ Plotted are the normalized abundance values for each of 3 replicates across the thermal profile with the best fit curves overlayed for each individual replicate. Similarly, the UQCRFS1 chaperone protein, LYR motif containing 7 (LYRM7), is also destabilized in the presence of SP-2577 (ΔTm = -3.0°C, Fig. 1D). Other members of Electron Transport Chain Complex III (ETCIII) are either not detected in our data (CYC1, QCR6-10), or show no significant change in Tm (QCR1 ΔTm = 0.01°C, QCR2 ΔTm = 0.1°C). Since ETCIII is a large transmembrane complex, it is likely not solubilized in the detergent-free CETSA-MS buffer. Supporting this, we found we were able to detect CYC1 when including detergent in the CETSA buffer, but were unable to detect changes in UQCRFS1 stability in these conditions (Supplemental Fig. 4,5). We thus posit that the ETCIII subunits detectable by CETSA-MS are not yet loaded into the full complex and are more vulnerable to destabilization by SP-2577.

To confirm the findings from the CETSA-MS experiment, we performed western blot coupled CETSA with SP-2509. Similar to our CETSA-MS, UQCRFS1 and LYRM7 both showed reduced stability with treatment (Fig. 2A). Interestingly, NADH:ubiquinone oxidoreductase core subunit S1 (NDUFS1), another mitochondrial protein from ETC Complex I (ETCI) was shown to be stabilized with treatment in the CETSA-MS assay (ΔTm = 1.2°C). However, we were instead surprised to find the CETSA Western blot showed that SP-2509 treatment also destabilized NDUFS1. Both UQCRFS1 and NDUFS1 contain Fe-S cluster cofactors, thus we investigated whether SP-2509 affects another key protein involved in the transfer of iron-sulfur clusters into nascent Fe-S-containing proteins, HSCB. ^17^

**Figure 2.**
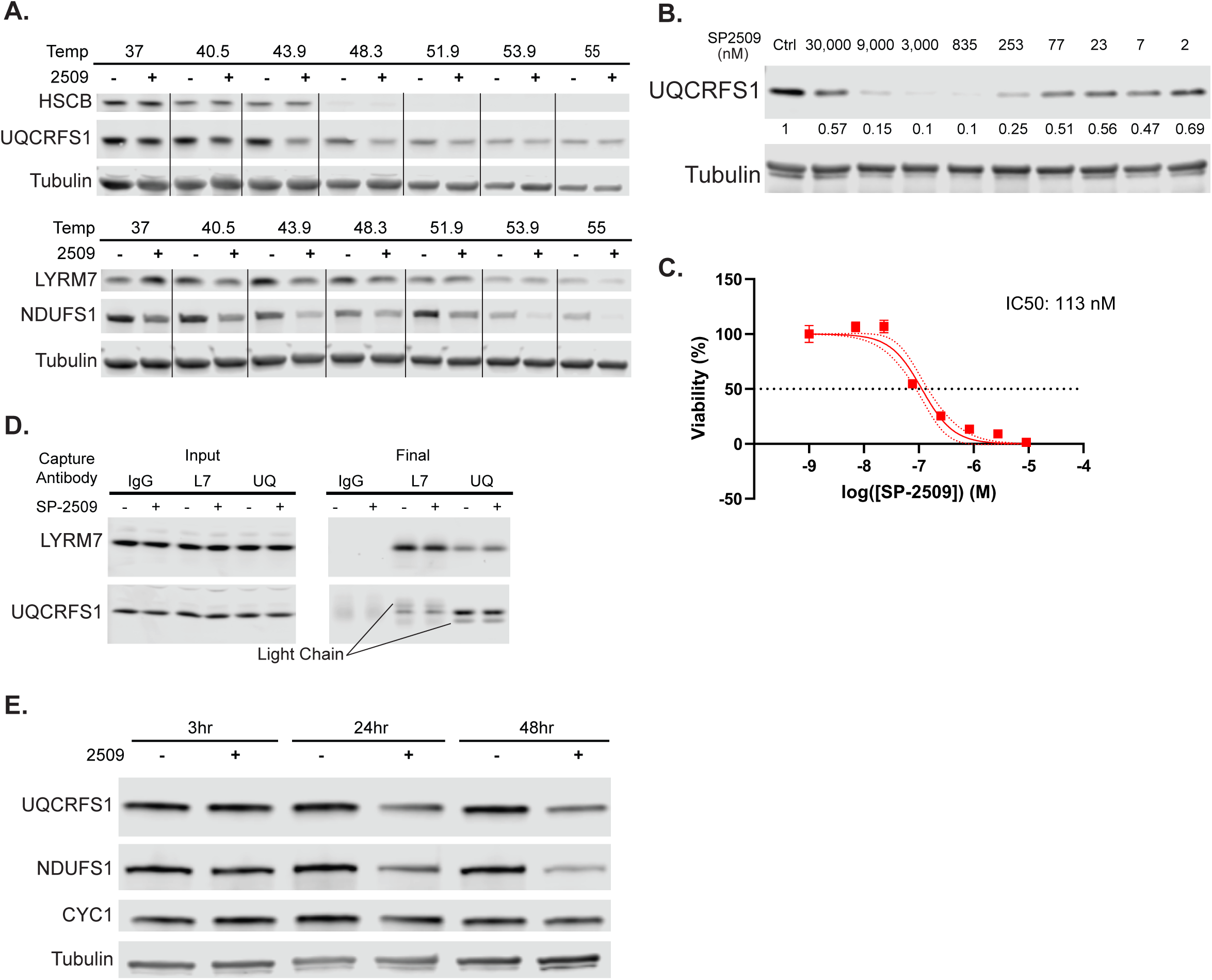
SP-2509 treatment destabilizes UQCRFS1 in a dose-dependent manner. All assays were performed in A673 cells. A) Cells were treated with either 0.1% DMSO or 1 µM SP-2509 for 3 hours and then soluble protein stability was analyzed by CETSA Western blot. B) Cells were treated with various doses of SP-2509 for 3 hrs and then soluble protein stability was analyzed by isothermal CETSA Western blot. Middle numbers represent tubulin normalized quantification relative to Ctrl using Licor Odessy software. C) Cells were treated with various doses of SP-2509 for 72 hours and then cell viability was assayed by Cell TiterGlo. Dots represent mean viability (N = 3), while the line is the curve of best fit using a 4 parameter logorithmic model. Error bars represent standard deviation, while dotted lines represent 95% confidence interval of the model. D) Cells were treated with 0.1% DMSO or 1 µM SP-2509 for 3 hours, after which UQCRFS1 and LYRM7 compexes were assayed by Co-IP. E) Cells were treated with 0.1% DMSO or 1 µM SP-2509 for 3, 24, or 48 hrs and then total protein content was assayed by Western blot.

### SP-2509 destabilizes UQCRFS1 at similar IC50 to cytotoxic activity

In biochemical assays with purified enzyme, SP-2509 inhibits LSD1 with an IC50 of ∼20 nM, but shows cytotoxicity in Ewing sarcoma cells at concentrations ranging from 100 nM – 2 µM *in vitro*. ^18^ The discordance between these observed IC50 values suggests that LSD1 inhibition with SP-2509 may not be the sole driver of cell death. We thus sought to determine the concentration of SP-2509 required to destabilize UQCRFS1. To do this, we performed an isothermal CETSA, treating samples with increasing concentrations of SP-2509 (2 – 30,000 nM) and heated to the Tm of UQCRFS1 (48 °C) (Fig. 2B). By plotting the quantified intensity of our western blots with respect to temperature we were able to show a dose-dependent destabilization of UQCRFS1, with over 50% destabilized above the observed cytotoxicity IC50 of 114 nM (Fig. 2C). This concordance between the concentrations that result in observed UQCRFS1 destabilization and cytotoxicity suggest that the mechanisms leading to impaired Fe-S cluster biogenesis may be linked to cell death in these cells.

### SP-2509 does not disrupt the LYRM7/UQCRFS1 complex

UQCRFS1 and LYRM7 destabilization could be caused by SP-2509 disrupting the ability of LYRM7 to complex with UQCRFS1 and act as a chaperone. To test this, we treated A673 cells with 1 µM of SP-2509 or DMSO and performed co-immunoprecipitation (Co-IP) using anti-LYRM7 and anti-UQCRFS1 antibodies. Our Co-IP showed no difference in the ability to pull down either protein during treatment with SP-2509 (Fig. 2D). This suggests that despite a precipitous drop in thermal stability upon SP-2509 treatment, UQCRFS1 and LYRM7 are likely able to form a stable complex.

### SP-2509 causes a time-dependent depletion of Fe-S cluster proteins

Another possible reason for the decreased stability of both UQCRFS1 and NDUFS1 could be absence of their Fe-S cofactors. Impaired incorporation of Fe-S clusters into proteins that require this cofactor has been shown to result in progressive protein depletion. ^19–21^ We assayed the total levels of UQCRFS1 and NDUFS1 following SP-2509 treatment for 3, 24, and 48 hrs (Fig. 2E). Both UQCRFS1 and NDUFS1 show marked reduction in total protein levels at 24 and 48 hrs. Conversely, cytochrome C1 (CYC1), a member of ETCIII with a heme cofactor, is not altered in the presence of SP-2509. These data suggest that SP-2509 specifically inhibits the proper folding of key Fe-S cluster proteins over this timescale, without disrupting a similar ETC protein containing a heme cofactor.

In human cells, Fe-S proteins constitute a relatively small fraction of the proteome, making proteome-wide analysis by mass spectrometry challenging. To address this, we concentrated our mass spectrometry analysis on the mitochondrial proteome, as many Fe-S cluster proteins localize to mitochondria. We then evaluated Fe-S protein abundance after 16 hours of treatment with either (1 µM) SP-2509 or DMSO by labeling tryptic digests of both cell lysates with reductive dimethylation using isotopic formaldehyde (ReDiMe) or tandem mass tags (TMT). In concordance with our previous CETSA-MS and CETSA western blots, both NDUFS1 and UQCRFS1 were depleted in SP-2509 treated cells. Additionally, the majority of other detected mitochondria-associated Fe-S proteins (∼75%) showed depletion following SP-2509 treatment (Fig. 3A,B Supplemental Table 2). FDX1, an essential component of steroidogenesis and lipid metabolism, and SDHB, which facilitates electron transfer from succinate to ubiquinone in the TCA cycle, were both most affected at this timepoint.

**Figure 3.**
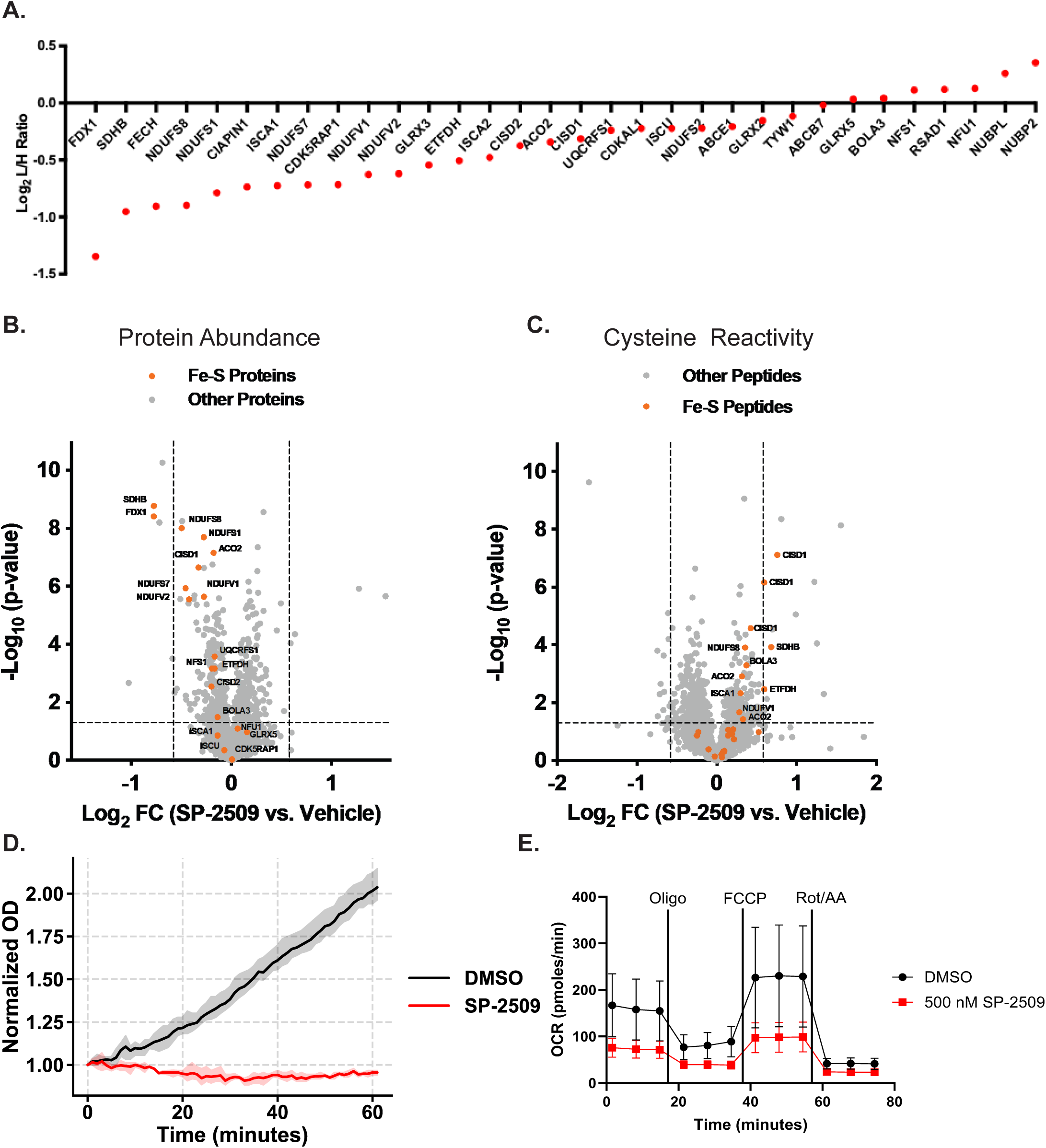
SP-2509 treatment broadly impacts abundance, cysteine reactivity, and activity of Fe-S cluster proteins in the mitochondria. All assays were performed in A673 cells. Cells were treated with 0.1% DMSO or 1 µM SP-2509 for 16 hrs and then protein content and cysteine reactivity were assayed by mass spectrometry. A) A plot of protein abundance of Fe-S containing proteins quantified by ReDiMe. B) A volcano plot of Log2(FC protein abundance) vs -Log10(p-value). Fe-S proteins are indicated in yellow and quantified by TMT. C) A volcano plot of Log2(FC cysteine reactivity) vs -Log10(p-value). Fe-S cysteine reactivity was quantified by desthiobiotin (DBIA) and TMT. D) Cells were treated with 0.1% DMSO or 1 µM SP-2509 for 24 hours and then Aconitase activity was assayed. Shaded region represents 95% confidence interval (N = 4) E) Cells were treated with either 0.1% DMSO or 500 nM SP-2509 for 24 hrs and then cellular respiration was assayed by Seahorse XF assay. Error bars represent standard deviation (N = 3).

The absence of Fe-S clusters exposes liganding cysteine residues and increases their reactivity towards cysteine-reactive probes such as desthiobiotin iodoacetamide (DBIA). We next assessed the relative reactivity of cysteine residues in peptides involved in Fe-S cluster binding using DBIA and TMT proteomics. Depleted mitochondrial Fe-S proteins show increased cysteine reactivity in the presence of SP-2509 (1 µM), indicating loss of Fe-S clusters in these proteins (Fig. 3C). Commensurate with the increase in cysteine reactivity, SP-2509 inhibits the activity of the Fe-S containing enzyme Aconitase, which acts as a transcription factor to regulate iron metabolism in during iron deprivation (Fig. 3D). ^22^ Thus, the reduced thermal stability of these Fe-S proteins is likely caused by impaired Fe-S cofactor binding and improper protein folding. Importantly, given the prevalence of Fe-S cluster proteins in the mitochondria, we were also able to observe impaired ETC function following prolonged SP-2509 exposure with decreases in both basal respiration and maximum respiratory capacity after 24 hrs of treatment (Fig. 3E).

### SP-2509 activity is antagonized by iron and copper supplementation

Fe-S cluster protein depletion was reported as a feature of cuproptosis. ^19^ In these experiments the copper ionophore, elesclomol, complexed with Cu^2+^ in the media and transported it across the cell membrane.^19^ In the cell Cu^2+^ is reduced to Cu^1+^, and Cu^1+^ can compete for binding of the sulfhydrl groups of Fe-S proteins with high affinity.^23^ Complexation with transition metals, including Cu^2+^, has also been proposed as mechanism for SP-2509 cytotoxicity^8^ and we wondered whether SP-2509 cytotoxicity involved copper as shown for elesclomol. We therefore tested whether addition of Cu2+ to the media potentiates SP-2509 cytotoxic activity and, conversely, whether the addition of the copper chelator tetrathiomolybdate (TTM) antagonizes SP-2509 activity, as was reported for elesclomol. ^21^ We used SynergyFinder to determine the degree to synergy and antagonism between SP-2509 and other treatments. A ZIP-score is a percent deviation from the expected measurement if there is no interaction. ^24, 25^ A score of 10 generally indicates synergy while a score of -10 generally indicates antagonism. Interestingly, TTM shows minimal antagonism with a min ZIP-score of -8.82, (Fig. 4A,B) while the addition of CuCl2 to SP-2509 treatment shows a strong antagonistic effect, with a min ZIP-score of -30.55, (Fig. 4D,E). These data suggest that SP-2509 is not a copper ionophore and does not cause cell death through canonical cuproptosis.

**Figure 4.**
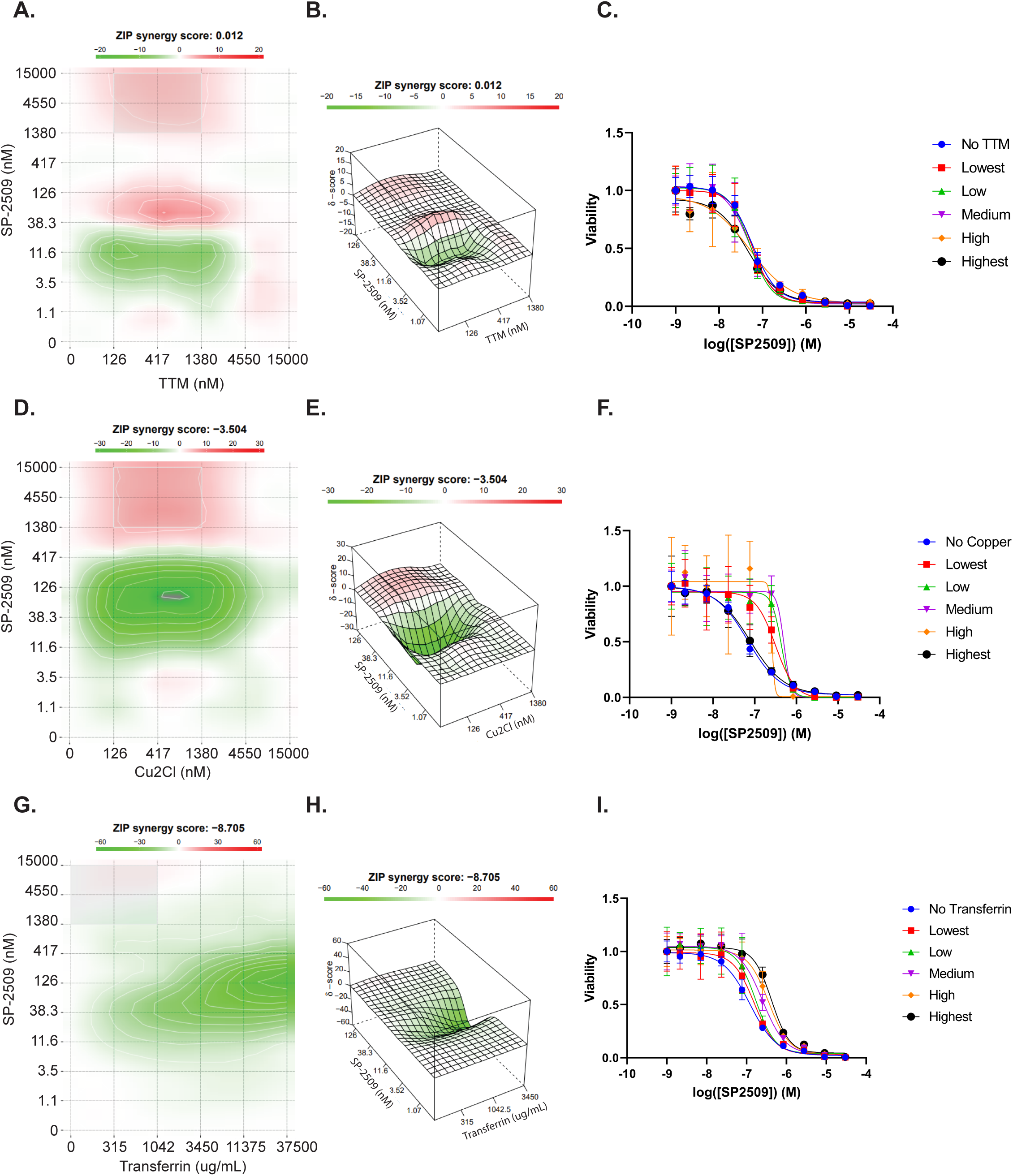
Transferring confers resistance to SP-2509 treatment. Cells were treated with various doses of SP-2509 and a secondary compound (Tetrathiol Molybdate, Cu2Cl, Transferrin, N = 3) for 72 hrs and then cell viability was assayed by Cell TiterGlo assay. A, B, D, E, G, H) Synergistic interactions were identified using Synergy finder software. C, F, I) Assays were also graphed as separate curves for each dose of secondary compound. Error bars represent standard deviation, while the lines represent the curve of best fit using a 4 parameter logorithmic model.

We next hypothesized that SP-2509 might instead disrupt the availability of iron for Fe-S cluster biogenesis. Direct supplementation of Fe(II) and Fe(III) is acutely toxic to cells at relevant concentrations. Instead, we supplemented SP-2509 treated cells with Transferrin, which is a key protein responsible for the cellular import of iron. ^26^ Transferrin supplementation exhibited an antagonistic effect (min ZIP-score = -49.38, Fig 4G,H), indicating that iron supplementation attenuates SP-2509 cytotoxicity. That both copper and iron supplementation reduce the potency of SP-2509 indicates a potential interaction between SP-2509 and these transition metals.

### SP-2509 cytotoxic activity is LSD1 independent

Though our data suggest real effects on the stability of Fe-S cluster proteins following treatment with SP-2509, the relationship between UQCRFS1 (and other Fe-S cluster protein) destabilization, LSD1 inhibition, and cytotoxicity was unclear. Because SP-2509 was identified as a non-competitive inhibitor, it has been challenging to determine whether cytotoxicity in Ewing sarcoma is the result of downstream effects from on-target LSD1 binding or other off-target effects. Moreover, as an epigenetic regulator, LSD1 can affect various aspects of mitochondrial metabolism in various cell contexts. In order to test whether cytotoxicity and UQCRFS1 destabilization were related to LSD1 inhibition, we used several additional tool compounds characterized in the original screen that identified SP-2509. ^18^ Several additional compounds (compounds 15, 19, and 20, Fig. 5A) were reported that had similar cytotoxic effects in the T-47D breast cancer cell line, but lacked detectable LSD1 inhibition in biochemical assays.^18^ Notably, these compounds possess the same N’-(2-hydroxybenzylidene)hydrazide core as SP-2509 and SP-2577. We tested the cytotoxicity of all 4 N’-(2-hydroxybenzylidene)hydrazide compounds in Ewing sarcoma cells and found them to have comparable potency at 72hrs (IC50s SP-2509: 110 nM, Cmpd 15: 124 nM, Cmpd 19: 166 nM, Cmpd 20: 170 nM, Fig 5B.). These results indicate that cytotoxic activity in Ewing sarcoma is likely LSD1 independent.

**Figure 5.**
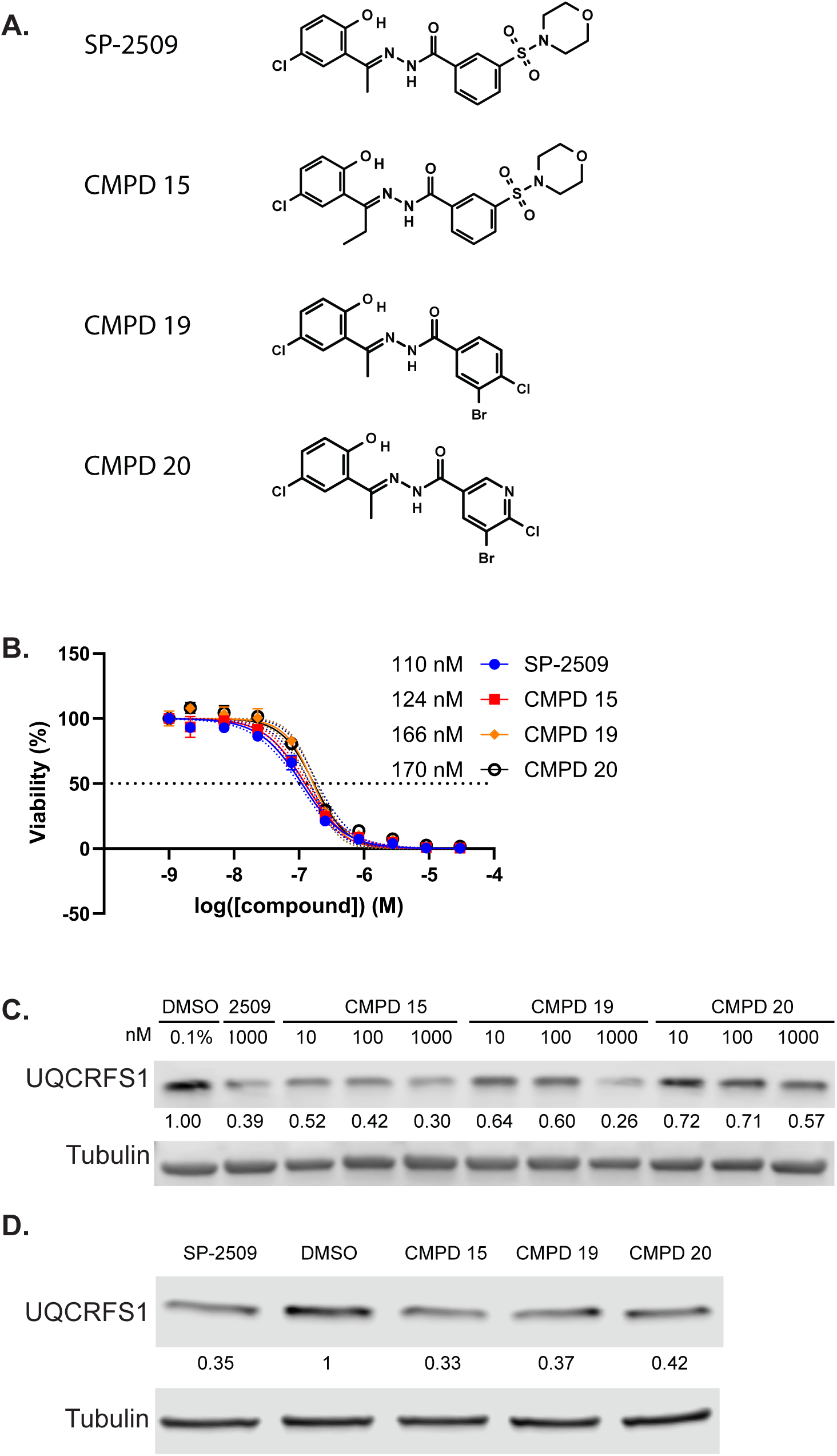
Other N’-(2-hydroxybenzylidene)hydrazides have similar activity in cytotoxicity and UQCRFS1 destabilization assays. A) Chemical structure of SP-2509 as well as compounds 15, 19, and 20. B) Cells were treated with various doses of SP-2509 or compounds 15, 19, or 20 for 72 hrs and then cell viability was assayed by Cell TiterGlo. C) Cells were treated with 0.1% DMSO, 1000 nM SP-2509 or various doses of compounds 15, 19, or 20 for 3 hrs and then soluble protein stability was assayed by isothermal CETSA Western blot. D) Cells were treated with 0.1% DMSO or 1 µM of SP-2509 or compound 15, 19, or 20 for 24 hours and then total protein content was assayed by Western blot. Middle numbers represent tubulin normalized quantification relative to Ctrl using Licor Odessy software.

Given that our data above suggest a mechanistic role for of SP-2509 in disrupting Fe-S cluster biogenesis and destabilizing Fe-S proteins we next asked whether this was also a function of the N’-(2-hydroxybenzylidene)hydrazide core. Compounds 15, 19, and 20 were assayed for their effects on UQCRFS1 stability at 3 hrs and abundance at 48 hrs. We found all three compounds destabilize UQCRFS1 at similar doses to SP-2509 (Fig. 5C) and cause similar depletion of UQCRFS1 protein at 48hrs (Fig. 5D).

Importantly, these data indicate that effects on Fe-S cluster biogenesis and the stability of Fe-S cluster proteins are also LSD1-independent. Instead, the N’-(2-hydroxybenzylidene)hydrazide core is linked to both Fe-S cluster disruption and cytotoxicity.

### Non-LSD1 inhibitory compounds recapitulate SP-2509 gene expression changes

SP-2509 treatment was previously shown to reverse the transcriptional activity of the oncogenic fusion transcription factor EWSR1::FLI1. Having linked Ewing sarcoma cell death to off-target disruption of Fe-S proteins using compounds 15, 19, and 20, we next asked whether the transcriptional activity seen in Ewing sarcoma cells was also associated with the N’-(2-hydroxybenzylidene)hydrazide core or on-target LSD1 inhibition. To test this, we treated A673 cells with 2 µM of drug (compound 15, 19, 20 or OG-L002) or vehicle control (DMSO) for 48 hrs and analyzed the transcriptome by RNA-seq. OG-L002 was used as a positive control for LSD1 inhibition without the N’-(2-hydroxybenzylidene)hydrazide core. These data were analyzed alongside previously published data of A673 cells treated with either 2 µM SP-2509 (SP-2509-b1) or DMSO (DMSO-b1) for 48 hrs, as well as A673 samples treated with shRNA targeting either EWSR1::FLI1 (iEF) or Luciferase (iLuc) as control.^5, 27^

Principal component analysis shows strong overlap between DMSO, OG-L002, and to a lesser extent iLuc samples (Fig. 6A). Samples treated with SP-2509, 15, 19, and 20 also showed strong overlap, except for a single replicate outlier for compound 20 (Fig. 6A). These samples shared a first principal component with iEF samples, but differed along the second component, indicating that while iEF and SP-2509 treatment share a dominant component of the gene expression changes, there are aspects unique to the EWSR1::FLI1 knockdown (Fig. 6A). This is consistent with the prior observation that SP-2509 treatment blocks EWSR1::FLI1 transcriptional activity. Differentially expressed genes (FDR < 0.05 & |FC| > 1.5) were plotted as batch corrected read counts, normalized by gene across samples (Fig. 6B). Hierarchical clustering allows for visual identification of 5 clusters of genes: those expressed predominantly after iEF treatment, those shared between SP-2509 treatment and iLuc, those shared between SP-2509 treatment and iEF, a cluster of genes shared between DMSO, OG-L002 and iEF samples, and finally those shared between all the control samples (DMSO and iLuc) and the OG-L002 samples.

**Figure 6.**
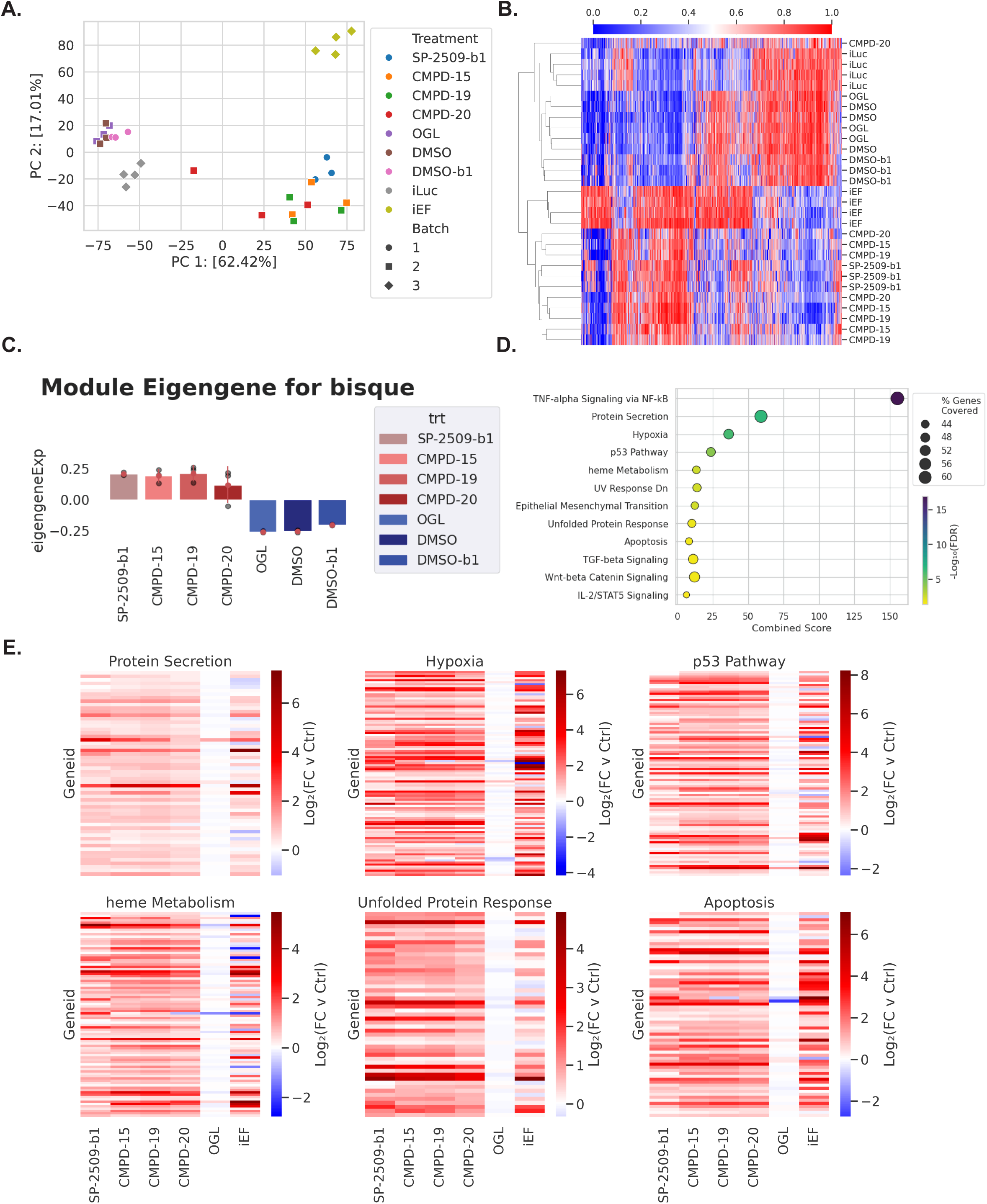
N’-(2-hydroxybenzylidene)hydrazides without LSD1 inhibitory activity block EWSR1::FLI1 transcriptional activity in cells. Cells were treated with 0.1% DMSO or 2 µM of OG-L002, or compound 15, 19, or 20 for 48 hrs and then transcriptome was analyzed by RNA-seq. This data was merged and batch corrected with previous data using pyComBat-seq. In previous data cells were treated with 0.1% DMSO (DMSO-b1), 2 µM SP-2509 (SP-2509-b1), or transduced with Luciferase (iLuc) or EWS::FLI1 (iEF) shRNAs for 48 hrs. A) A graph of principle component analysis using principal component 1 vs principal component 2. B) Heatmap of differentially expressed genes (FDR < 0.05, |FC| > 1.5) as identified by pyDESeq. Batch corrected counts were normalized on a per gene basis. C) Expression of bisque Eigengene identified using pyWGCNA, which contains genes highly expressed in SP-2509 treated cells compared to control using batch corrected counts. D) GSEA analysis of genes contained in bisque module vs MSigDB 2020 gene sets (FDR < 0.05). E) Heatmaps of genes contained in select pathways identified in bisque module. Data is DESeq log 2 FC values vs control samples (DMSO and iLuc).

Weighted gene network connectivity analysis (WGCNA) was used to identify a cluster of genes that exhibit jointly increased expression in samples treated with SP-2509, 15, 19, and 20 as compared to DMSO and OG-L002. The shRNA samples were excluded from this analysis to better isolate the effects of N’-(2- hydroxybenzylidene)hydrazide treatment (Fig. 6C). The genes identified in this cluster were used for gene set enrichment analysis (GSEA) against the MSigDB Hallmark 2020 gene sets. GSEA identified several pathways whose gene sets were enriched in SP-2509 treated samples (Fig. 6D, FDR < 0.05). Upregulated pathways include protein secretion, hypoxia, and heme metabolism. The protein secretion pathway includes mechanisms necessary for cyclical endocytosis and secretion of Transferrin to import extracellular iron. ^28^ Heme metabolism and hypoxia pathways are linked to the iron content and oxygen carrying capacity of Heme. ^29, 30^ These pathways are co-upregulated suggesting engagement of iron dependent metabolic pathways and likely constitutes an iron dependent compensatory response. Additional upregulated pathways include the unfolded protein response (UPR), p53 pathway, and apoptosis. The instability of Fe-S containing proteins in response to SP-2509 treatment may explain the activation of UPR seen here and in previous studies. ^3^ Further, both the presence of UPR as well as the dearth of key mitochondrial Fe-S containing proteins could activate of p53 and apoptotic pathways that contribute to cell death. ^31, 32^

The constituent genes for each pathway were identified and heatmaps were generated for each treatment group compared to shared control samples using log 2-fold change values generated by DESeq2 (Fig. 6E). Gene expression in iEF samples shows considerable overlap with SP-2509 treated samples but is less concordant than samples treated with compounds 15, 19, or 20. Taken together, these data show that treatment of Ewing sarcoma cells with N’-(2-hydroxybenzylidene)hydrazide compounds leads to a reversal of oncogenic gene expression caused by EWSR1::FLI1. Given that compounds 15, 19, and 20 lack LSD1 inhibitory activity, these data indicate that the EWSR1::FLI1 targeting activity of these compounds is LSD1-independent and may be downstream from disruptions in Fe-S biogenesis. Further studies will be required to discern the biological mechanisms linking Fe-S cluster biogenesis to EWSR1::FLI1 biology, as well as to understand SP-2509 induced cell death and the mechanisms that drive resistance to SP-2509 treatment.

## DISCUSSION

The ability of SP-2509 and related compounds to block the transcriptional regulatory activity of the oncogenic fusion transcription factor EWSR1::FLI1 and impair tumor growth has led to the clinical development of SP-2577 for Ewing sarcoma and related tumors. However, despite much research into SP-2509 as an LSD1 inhibitor, the role for LSD1 in Ewing sarcoma and the mechanistic basis for N’-(2- hydroxybenzylidene)hydrazide-mediated cell death both remain poorly understood. Other LSD1 inhibitors lacking activity against Ewing sarcoma and screening for resistant phenotypes converged to suggest a mitochondrial role in SP-2509 activity. ^314^ These prior reports raised the possibility of either 1) a link between LSD1 and the mitochondria in Ewing sarcoma cells or 2) off-target activity involving the mitochondria. In this study, we use both genomic and chemical approaches to investigate the LSD1 dependence of SP-2509 activity, including cytotoxicity and anti-EWSR1::FLI1 transcriptional regulation. Further, our unbiased proteomics and validation studies link this activity to impaired Fe-S cluster biogenesis and destabilization of proteins requiring an Fe-S cofactor. A chemoproteomic evaluation of Fe-S proteins shows pleiotropic effects following SP-2509 treatment, and this may explain the relatively modest resistance shown after knocking down any single member of the electron transport chain in previous work. ^14^ The exception to this is MRPL45, a member of the mitochondrial ribosome, the removal of which inhibits the production of all translated proteins of the mitochondrial genome.

Previous studies of elesclomol and its role as a copper ionophore have shown similar depletion of Fe-S cluster proteins during cuproptosis, a copper mediated form of cell death. ^19^ While SP-2509 mediated cell death shows a similar depletion in Fe-S cluster proteins, supplementation with copper causes resistance rather than the increased potency reported for elesclomol. This suggests that, while SP-2509 derivatives may have some copper binding capacity, the mechanism of cell death is not copper dependent. This is further supported by the failure of the copper chelator TTM to substantially change the IC50 of SP-2509 (Fig. 3A). Considering these findings, it appears likely that SP-2509 derivatives are likely restricting the bioavailability of iron in the cell and that complexing with copper in the media may diminish this activity. The affinity of SP-2509 for individual metal ions is an active area of study, as are the features of iron metabolism in the presence of SP-2509 and related compounds.

Several gene signatures in our RNA-seq data support a role for SP-2509 in disrupting iron metabolism in the cell, including pathways involved in protein secretion, hypoxia, heme metabolism, and unfolded protein response. Altered iron metabolism could explain the increased expression of heme metabolism related genes, as well as engagement of protein secretion related genes and hypoxia related genes.^28–30, 33^ To extract iron from the extracellular environment, cells must endocytose Transferrin with iron bound to it. Transferrin then releases bound iron in the low pH environment of the lysosome before being secreted back out of the cell. The hypoxia response may be explained by increased expression of the iron responsive hypoxia transcription factor EPAS1, which shows substantially increased expression in response N’-(2- hydroxybenzylidene)hydrazide treatment.^34^ If these iron responsive pathways are unable to protect the cell from SP-2509 treatment the Fe-S cluster protein instability goes on to trigger UPR response genes and eventually cell death. This is consistent with our previous findings that SP-2509 treatment strongly induces the ER stress and UPR pathway after 24 hrs. ^3^ Additional studies are needed to further define these mechanistic details.

How disrupting Fe-S clusters leads to transcriptional changes seen in the RNA-seq data remains intriguing and is a topic of ongoing study. Importantly, compounds 15, 19, and 20 do not inhibit LSD1, but retain the ability to reverse the transcriptional signature of EWSR1::FLI1. This suggests that gene expression changes are downstream of the Fe-S cluster inhibition. Many possible mechanisms exist to explain this finding. Some aspects of EWSR1::FLI1 activity may depend on nuclear proteins that require iron or Fe-S cofactors. Alternatively, there may be some metabolic feedback between the mitochondria and nucleus. Another possibility is that the transcriptional similarities between N’-(2-hydroxybenzylidene)hydrazide treatment and EWSR1::FLI1 arise from convergence of purpose. In the same way that SP-2509 treatment deprives the cell of essential nutrients and the cell responds in a compensatory manner, the expression of EWSR1::FLI1 is required by the Ewing sarcoma cells to activate the nutrient uptake and growth pathways that allow for proliferation. The removal of this essential transcription factor similarly deprives the cell of nutrients and disrupts the translation of growth-related genes, thus activating a similar set of compensatory pathways. This could explain why prior reports found that EWSR1::FLI1 knockdown conferred greater resistance than that seen for knockdown of individual mitochondrial components. ^2^

Importantly, this study makes significant progress toward defining the mechanistic basis for SP-2509 and SP-2577 activity in Ewing sarcoma, and likely other cell lines. The ability of SP-2509 to potently inhibit the maturation of Fe-S cluster proteins raises interesting question regarding the utility of N’-(2- hydroxybenzylidene)hydrazides as anti-cancer agents. Iron metabolism and the maturation of Fe-S containing ETC complex members is not unique to cancer cells, however, early clinical trials of SP-2577 show a manageable safety profile with preliminary signs of activity albeit in a very small number of patients. ^35^ Additionally, our recent work found that SP-2509 and SP-2577 exhibits activity against a range of fusion driven sarcomas, but the determinants of activity against any particular cell of interest is a topic of future study. ^36^ Our data suggest that the anti-LSD1 activity of SP-2509 is completely ancillary, therefore future derivative compounds can be optimized toward improved pharmacokinetics, pharmacodynamics, and physicochemical properties without being restricted by the ability to bind and inhibit LSD1. Ultimately, better understanding of the pharmacology of these compounds and how they lead to cell death will enable more intelligent clinical translation moving forward.

## METHODS

### Reagents

SP-2509 and OG-L002 were purchased from Selleck Chemicals (Cat. # S7680 and S7237, respectively.) Seclidemstat was provided by Salarius Pharmaceuticals. Compounds 15, 19, and 20 were synthesized by the Medicinal Chemistry Shared Resource at The Ohio State University James Comprehensive Cancer Center. Identity and purity were verified by HPLC-MS and NMR.

### Antibodies

The following antibodies were used for immunodetection: anti-α-tubulin (DM1A; Abcam ab7291), IRDye® 800CW goat anti-mouse IgG (LI-COR Biosciences 926-32210), IRDye® 800CW goat anti-rabbit IgG (LI-COR Biosciences 926-32211), anti-UQCRFS1 (Abcam ab191078), anti-LYRM7 (Novus NBP2-14701), anti-NDUFS1 (Abcam ab169540), anti-CYC1 (Abcam ab137757).

### CETSA-Mass Spectrometry Sample Preparation

A673 cells were treated for 3 hrs with SP-2577 (Seclidemstat, 1 µM), an SP-2509 derivative, or DMSO (0.1%), and then divided evenly into 5 samples. Samples were then heated to 37, 47, 52, 57, and 62 °C respectively for 5 min, resulting in irreversible protein unfolding proportional to temperature. Soluble proteins were separated from insoluble unfolded, and membrane bound proteins by centrifugation, and then analyzed by proteomic mass spectrometry at the OSU CCIC Proteomics core. Briefly, samples were digested with trypsin, concentration of tryptic peptides were measured, equal amount of the peptides were then individually labeled with different TMT mass labels following manufacture’s instruction. Labeled samples were then mixed and fractionated to enable multiplexing and quantitation of the samples for each replicate.

### CETSA-MS Analysis

Protein detection quantities were normalized to the 37 °C samples to generate fold-change values. All curve fitting was generated by first attempting the Levenberg-Marquardt method, if that was unsuccessful gradient descent was used instead. Curves were first fit to the median values for each replicate. Theses curves were then used to generate correction factors for each replicate such that all curves were shifted to overlap with the best fitting replicate. Individual proteins that failed to melt, or conversely reached 0.25 fold change by 47 °C, were filtered. The correction factors calculated for each replicate were then applied to all respective proteins before curve fitting was attempted for each protein. The IC50 is defined as the inflection point of the resulting curve. Curves were excluded from further analysis if they met any of the following criteria:

- where the inter-replicate standard deviation (IRSD) in IC50 is outside 2 standard deviations of the IRSD across all proteins
- where the reduced mean squared error of the fit is greater than 0.2
- where the calculated mean IC50 in is greater than 62 °C

The resulting curves and corrected fold-change values were then processed and graphed using python. Graphs were generated using seaborn and matplotlib.

### CETSA Sample Preparation

Cells were seeded to be 90-100% confluent in a 15 cm dish by the following day. Cells were then dosed with treatment or control media and incubated for 3 hrs. 8 mL of media from each plate was saved, cells were trypsinized and the reaction was quenched with the saved media. Each sample was pelleted by centrifugation at 300xg for 5 min and then resuspended in 400 µL of HBSS + protease inhibitor at approximately 2x10^6^ cells per 50 µL. Each sample was divided into 50 µL increments and aliquoted into the PCR strip tubes and immediately heated for 5 min using the gradient function of a PCR machine. Following heating, the samples were immediately lysed by 3 rounds of freezing in liquid nitrogen followed by thawing on the benchtop at room temperature for 5 min. After the final round of freeze thaw, samples were transferred to 1.5 mL centrifuge tubes and centrifuged at 12,000xg for 15 min at 4 °C to clear the lysate. The supernatants of each sample were transferred to a fresh set of PCR strip tubes, mixed with Laemmli buffer and boiled for 5 min to prepare the samples for SDS-PAGE.

### Isothermal CETSA Sample Preparation

Cells were seeded to be 90-100% confluent in 6-well dishes by the following day. Cells were then dosed with either vehicle control or various concentrations of each treatment and then incubated for 3 hours. 1 mL of media was saved from each well and then used for quenching after those wells had been trypsinized. Each sample was transferred to a 1.5 mL centrifuge tube and centrifuged at 300xg for 5 min and then resuspended in 30 µL of HBSS + protease inhibitor. All samples were then heated at 48 °C for 5 min. Samples were then prepared as described above in 1.5 mL centrifuge tubes and were thawed at room temperature in a metal heating/cooling block for 5 min after each freeze-thaw cycle.

### Whole Cell Protein Extraction

Approximately 1x10^6^ cells were trypsinized and then washed with HBSS before being transferred into 30 µL of RIPA buffer + 1:500 protease inhibitor cocktail (Sigma-Aldrich: P8340). Cells were lysed by pipetting up and down and the homogenized by passing the lysate through an insulin syringe. Lysate was then clarified by centrifugation at 12,000xg for 15 min at 4 °C. Total protein concentration was assessed via BCA assay (Thermo Scientific cat: 23225) according to manufacturer’s instructions. Samples were then diluted with additional RIPA buffer + protease inhibitor to have matching concentrations, mixed with Laemmli buffer and boiled for 5 min to prepare for SDS-PAGE.

### Immunoprecipitation

Cells were treated with 0.1% DMSO or 1 µM SP-2509 for 3 hours. Dynabead Protein A magnetic beads were precoated with 3% BSA in HBSS + 1 mM DTT. 1x10^7^ cells were trypsinized and resuspended in 550 µL of HBSS + protease inhibitor + 1 mM DTT. Samples were lysed and clarified by 3x freeze thaw as described for Isothermal CETSA. BCA assay was used to quantify protein concentration and samples were diluted to 2 µg/ml total protein. Lysate was incubated with 20 µL of beads to clear nonspecifically bound proteins for 1 hour at 4 °C. Samples were then incubated with 1 µg IP antibody rocking overnight at 4 °C. 50 µL of beads were added to each sample and incubated at 4 °C rocking for 2 hours. Beads were then washed 3x with ETNIP140 buffer at 4 °C rocking for 2 min. Protein was eluted from beads by boiling for 5 min in 1x Laemmli buffer diluted into RIPA buffer.

### Western Blot

20 µg of total protein was loaded into a BIORAD 4-15% Mini-PROTEAN TGX precast polyacrylamide gel and run at 120 V for 55 min using BIORAD Tris/Glycine/SDS electrophoresis buffer (25mM Tris, 192 mM glycine, 0.1% SDS, pH 8.3). Protein was transferred to Millipore Immobilon-FL PVDF membrane (0.45 µM pore size) by wet transfer using Towbin Buffer (diluted with methanol to working concentration the day of) at constant current 200 mA for 2 hours. Membranes were then blocked for 30 min in Licor Intercept (PBS) Blocking buffer, before being transferred to primary antibody incubation with antibody diluent (Intercept blocking buffer + 0.05% Tween 20) + 1:1000 dilution of primary antibody. Primary antibody incubation took place rocking, overnight at 4 °C. Membranes were then washed 3x in TBST rocking at RT for 5 min and incubated with appropriate Licor IRDye secondary antibody, diluted 1:12000 in antibody diluent, for 1 hour at room temperature while rocking. Excess secondary was washed once in TBST and twice in TBST + 0.01% SDS for 5 min at RT. Blots were resolved and band quantitation was performed using Licor Odyssey CLx imaging system.

### Cell Viability Assays

96-well plates were seeded with 2,500 cells per well and allowed to grow overnight. The following day treatment was added and cells were allowed to grow for 72 hours. All treatments and DMSO control were applied in triplicate. After 72 hours, cell viability was measured as total intra-cellular ATP content using the Cell Titer Glo (Promega) according to manufacturer instructions and data were analyzed using Graphpad Prism. Nonlinear regression was performed with a 4-point dose response curve. 2D cell titer glo was used to detect synergy and data were analyzed using SynergyFinder 3.0.

### RNA-sequencing

A673 cells were treated with 2 µM compound of interest for 48 hrs in triplicate. After 48 hours, cells were harvested, and RNA was extracted using RNeasy kit (Qiagen). RNA concentration was assayed by Nanodrop, and samples were submitted to the Institute for Genomic Medicine (IGM; Abigail Wexner Research Institute at Nationwide Children’s Hospital, Columbus, OH) following IGM sample submission guidelines. Samples were sequenced using a Hi-Seq 4000 to generate 150 bp paired-end reads. Reads were aligned using STAR aligner (2.7.9a) to the hg38 human genome. Differential gene expression was determined using pyDEseq ^37, 38^ while co-expressed gene modules were identified by first batch correcting normalized counts using pyCombat-seq ^39, 40^ followed by pyWGCNA ^41, 42^. Gene set enrichment analysis was performed on gene modules against the MsigDB Hallmark 2020 gene sets. Graphs were generated using the seaborn and matplotlib packages.

### Aconitase assay

Enzymatic activity of aconitase was measured using a commercially available kit from Cayman Chemical (Item no. 705502) according to manufacturer instructions with the modification that activity was measured for 1 hour rather than 30 minutes. Samples were assayed in triplicate and blank measurements for each timepoint were subtracted from sample measurements.

### Seahorse XFp assays

Metabolic flux analysis was performed using a Seahorse XFp flux analyzer (Agilent) following the manufacturer’s protocols. Cells were treated with either 0.1% DMSO or 500 nM SP-2509 for 24 hours prior to Seahorse XFp analysis. For mitochondrial stress tests, inhibitors (oligomycin, FCCP, rotenone/antimycin A) were prepared according to the manufacturer’s protocol (Agilent). Data were analyzed using GraphPad Prism.

### Crude Mitochondria Isolation

Crude mitochondria were obtained following a procedure adapted from previous literature. ^43^ A673 cells were collected in ice-cold PBS and washed twice with mitochondrial isolation buffer (IBc: 10 mM Tris/MOPS, 1 mM EGTA/Tris, 200 mM sucrose, pH 7.4). Cells were Potter-Elvehjem homogenized in IBc and homogenate was subjected to differential centrifugation. Cell debris and nuclei were pelleted at 600 g, 4 °C for 10 minutes and mitochondria were obtained by centrifuging the supernatant at 10,000 g, 4 °C for 10 minutes. Crude mitochondrial pellets were washed with PBS three times and protein concentration was quantified by DC Protein Assay Kit (Bio-Rad) before storage at -80 °C.

### Reductive Dimethylation (ReDiMe) of A673 Mitochondria for Protein Abundance Quantification (Figure. 3A)

A673 cells were treated with 0.1% DMSO or 1 µM SP-2509 for 16 hrs and mitochondria were isolated. 100 µg of each proteome was precipitated by the addition of 5 µL of 100% trichloroacetic acid/PBS and frozen. Upon thawing, protein pellets were obtained by centrifugation at 16,000 g, 4 °C for 10 minutes and washed with ice-cold acetone once. Pellets were then resuspended in 30 µL of 8 M urea/100 mM TEAB and the solution was diluted to 100 µL with 100 mM TEAB. Proteins were reduced with 1.5 µL of 1 M DTT (65 °C, 15 mins) and alkylated with 2.5 µL of 0.4 M iodoacetamide (37 °C, 30 mins). Reactions were diluted with 100 mM TEAB (120 µL) and incubated overnight with 2 μg of sequence-grade trypsin (Promega) and 1 mM CaCl2 at 37 °C. The digested proteomes were reductively dimethylated with 4 µL of 20% light (SP-2509) or heavy (DMSO) formaldehyde and 20 µL of 0.6 M sodium cyanoborohydride for 2 hours. Reactions were quenched by ammonium hydroxide, combined and desalted with a Sep-Pak C18 cartridge (Waters). Eluted peptides were dried in a SpeedVac chamber and resuspended in 400 µL of high-pH buffer A (95% water, 5% acetonitrile and 10 mM ammonium bicarbonate). Off-line fractionation was performed on a 25-cm Agilent Extend C18 column with a 60-minute linear gradient of 20% to 35% high-pH buffer B (10% water, 90% acetonitrile and 10 mM ammonium bicarbonate) and the six pooled fractions were dried by SpeedVac.

Peptides were resuspended in low-pH buffer A (100% water, 0.1% formic acid) and subjected to mass spectrometry analysis.

### Tryptic Digestion and TMT Labeling of A673 Mitochondria for Protein Abundance Quantification (Figure. 3B)

100 µg of mitochondria (n=3) from A673 cells treated with either 0.1% DMSO or 1 µM SP-2509 for 16 hrs were lysed in 100 µL of PBS. Proteins were denatured with 8 M urea/50 mM TEAB (pH 8.5), reduced by 10 mM DTT (65 °C, 15 mins) and alkylated with 25 mM iodoacetamide (37 °C, 30 mins). Proteins were precipitated by 600 µL of methanol, 200 µL of chloroform and 100 µL of water and centrifuged at 16,000 g, 4 °C for 10 minutes. Protein pellet was washed with ice-cold methanol once and resuspended in 300 µL of 0.2 M EPPS buffer (Thermo Scientific). Samples were digested overnight at 37 °C by 2 μg of sequence-grade trypsin (Promega) in the presence of 1 mM CaCl2. 30 µL of the digested peptides were diluted with 5 µL of EPPS buffer and 10 µL of acetonitrile prior to labeling by TMTsixplex isobaric tags (20 µg/µL, Thermo Scientific) for 60 minutes at room temperature (channels 126, 127, 128: DMSO; channels 129, 130, 131: SP-2509). Reactions were quenched with 5 µL of 5% hydroxylamine (Thermo Scientific) for 15 minutes and acidified with 5 µL of formic acid (Millipore Sigma). All channels were combined and dried in a SpeedVac chamber and desalted using Sep-Pak C18 cartridges (Waters).

### Cysteine Reactivity Profiling Using Desthiobiotin Iodoacetamide (Figure. 3C)

500 µg of mitochondria (n=3) from A673 cells treated with either 0.1% DMSO or 1 µM SP-2509 for 16 hrs were lysed in 500 µL of PBS. Proteome solutions were treated with 5 µL of 10 mM desthiobiotin iodoacetamide in DMSO for 1 hour at room temperature. Proteins were precipitated with methanol/chloroform as described above and resuspended in 9 M urea/50 mM TEAB (pH 8.5). Samples were reduced by 10 mM DTT (65 °C, 15 mins) and alkylated with 50 mM iodoacetamide (37 °C, 30 mins). Reactions were diluted with 300 µL of 50 mM TEAB and digested with 2 µg of trypsin overnight at 37°C in the presence of 1mM CaCl2. 50 µL of washed streptavidin agarose resin (Thermo Scientific) were added to each sample and rotated for 2 hours at room temperature. Upon removing the supernatant, the beads were washed with 3 x 1 mL of wash buffer (50 mM TEAB pH 8.5, 150 mM NaCl, 0.2% NP-40), 3 x 1 mL of PBS and 3 x 1 mL of water. Peptides were eluted with 50% acetonitrile/0.1% formic acid and washed twice more with acetonitrile. The combined supernatant was dried by SpeedVac. Peptides were labeled by TMT reagents (channels 126, 127, 128: DMSO; channels 129, 130, 131: SP-2509) and desalted as described above.

### Liquid Chromatography-Tandem Mass Spectrometry (LC-MS/MS) and Proteomic Data Analysis

LC-MS/MS was performed on an Orbitrap Exploris 240 Mass Spectrometer running Xcalibur v4.4 (Thermo Fisher Scientific) coupled to a Dionex Ultimate 3000 RSLCnano system. 5 µL of sample were loaded onto an Acclaim PepMap 100 C18 Loading Column (Thermo Scientific), followed by elution onto an Acclaim PepMap RSLC Column (Thermo Scientific). Peptides were separated by a 2-hour gradient of 5 to 25% Buffer B (20% water, 80% acetonitrile, 0.1% formic acid) in Buffer A at a flow rate of 0.3 µL/minute. The spray voltage was set to 2.1 kV. One full MS1 scan (120,000 resolution, 350-1,800 m/z, RF lens 65%, AGC target 300%, automatic maximum injection time, profile mode) was obtained every 2 s, with dynamic exclusion (repeat count 2, duration 10 s), isotopic exclusion (assigned) and apex detection (30% desired apex window) enabled. Varying numbers of MS2 scans (15,000 resolution, AGC target 75%, maximum injection time 100 ms, centroid mode) were obtained between each MS1 scan based on the highest precursor masses and filtered for monoisotopic peak determination, theoretical precursor isotopic envelope fit, intensity (5E4) and charge state (2-6). MS2 analysis included the isolation of precursor ions (isolation window 2 m/z) and HCD (collision energy 30%). LC-MS/MS data was analyzed in Thermo Proteome Discoverer v2.4 and searched against the Homo sapiens proteome database (SwissProt) using SequestHT and Percolator^1^. Trypsin was specified as the protease and maximum number of missed cleavages was set to 2. Peptide precursor mass tolerance was set to 10 ppm with a fragment mass tolerance of 0.02 Da. Oxidation of methionine (+15.995) and acetylation (+42.011) and/or loss of methionine (-131.040) of the protein N-terminus were set as dynamic modifications. For ReDiMe analysis, cysteine alkylation (+57.021) and either light- (+28.031) or heavy- (+34.063) dimethylation of lysine residues and peptide N-terminus were set as static modifications. For protein abundance TMT analysis, cysteine alkylation (+57.021) and TMT6plex modification (+229.163) of lysine residues and peptide N-terminus were set as static modifications. For cysteine reactivity quantification, carbamidomethyl (+57.021) and desthiobiotin iodoacetamide (+296.185 Da) were set as a dynamic modifications on cysteines and TMT6plex (+229.163) as static modifications on lysine residues and peptide N-termini. The FDR was set to 1% for peptide identification. Ratios generated were normalized by their respective channel medians. For TMT analysis, ratios were generated by dividing each channel with channel 126 and the average values of all ratios for each treatment condition from both biological replicates were calculated. Cysteine reactivity data was normalized with protein abundance data to yield net cysteine reactivity changes. Data was plotted in Prism 10 (GraphPad) and statistical analysis was performed using two-tailed t-test and Fisher’s combined probability test.

### Statistical Analysis

Dose response curves for cell viability data were determined in GraphPad Prism 9 using the 4-parameter, variable slope log(inhibitor) vs. response equation. 2-D dose response curves were determined using Synergyfinder 3.0. Three technical replicates were included in each of two biological replicates. Differentially expressed genes in the RNA-seq data were defined as genes with a multiple hypothesis testing adjusted (FDR/Benjamini-Hochberg^44^) p-value of <0.05 and an absolute log2 fold-change of >1.5. The distribution of the CETSA-MS data has excessive kurtosis which results in the data being non-normally distributed and the resulting p-values should be interpreted with care. P-values were determined by permutation test of means (scipy stats package, N = 100,000) followed by Benjamini-Hochberg correction, this emphasizes proteins with greater changes in IC50 while ignoring high standard deviation associated with this data. Proteins with p-values < 0.05 were considered worth further consideration.

### Data Availability

RNA-Seq data can be found in NCBI GEO with accession number GSE286100.

## ACKNOWLEDGEMENTS

We thank Ali Snedden, Yuan Zhang, and John Burian and the High-Performance Computing group at Nationwide Children’s Hospital for their support. We also acknowledge resources from the OSU Comprehensive Cancer Center (OSUCCC) Medicinal Chemistry Shared Resource (MCSR). We are grateful to the members of the Theisen Lab for comments and discussion of this manuscript during its preparation. This research was supported by institutional startup funds awarded to E.R.T., research project funds awarded by Salarius Pharmaceuticals to E.R.T., the American Cancer Society Research Scholar Grant RSG-22-118-01-DMC awarded to E.R.T., NIH T32CA269052 to J.S, and R35GM134964 to E.W.

