## Supplementary Figures for "N’-(1-phenylethylidene)-benzohydrazide cytotoxicity is LSD1 independent and linked to Fe-S cluster disruption in Ewing sarcoma"

### SUPPLEMENTARY MATERIALS

#### SUPPLEMENTARY TABLES

**Supplementary Table 1.** Melt curve fit data from derived from CETSA-mass spec data.

**Supplementary Table 2.** Proteomic data related to Main Figure 3.

**Supplementary Table 3.** Results from differential gene expression analysis of RNA-seq data

#### SUPPLEMENTARY FIGURES

**Supplementary Figure 1.** Histogram of individual ΔTm values (3 ΔTm values per protein, 1 per replicate).

**Supplementary Figure 2.** (A-D). Additional proteins with significant ΔTm's. On the top graphs, dots show corrected detectable fraction (N = 3), lines show the line of best fit for each replicate (N = 3). On bottom graphs, hash marks show the mean inflection point from the line of best fit. The dashed line indicates the 95% confidence interval for the inflection point.

**Supplementary Figure 3.** CETSA Western blot for MTRO4 (predicted band size ~ 31 kDa)

**Supplementary Figure 4.** CETSA Western blot for UQCRFS1 (top) and CYC1 (bottom) without NP-40.

**Supplementary Figure 5.** CETSA western blot for UQCRFS1 (top) and CYC1 (bottom) with 0.2% NP-40 in the HBSS used for protein extraction.

**Supplementary Figure 1.**

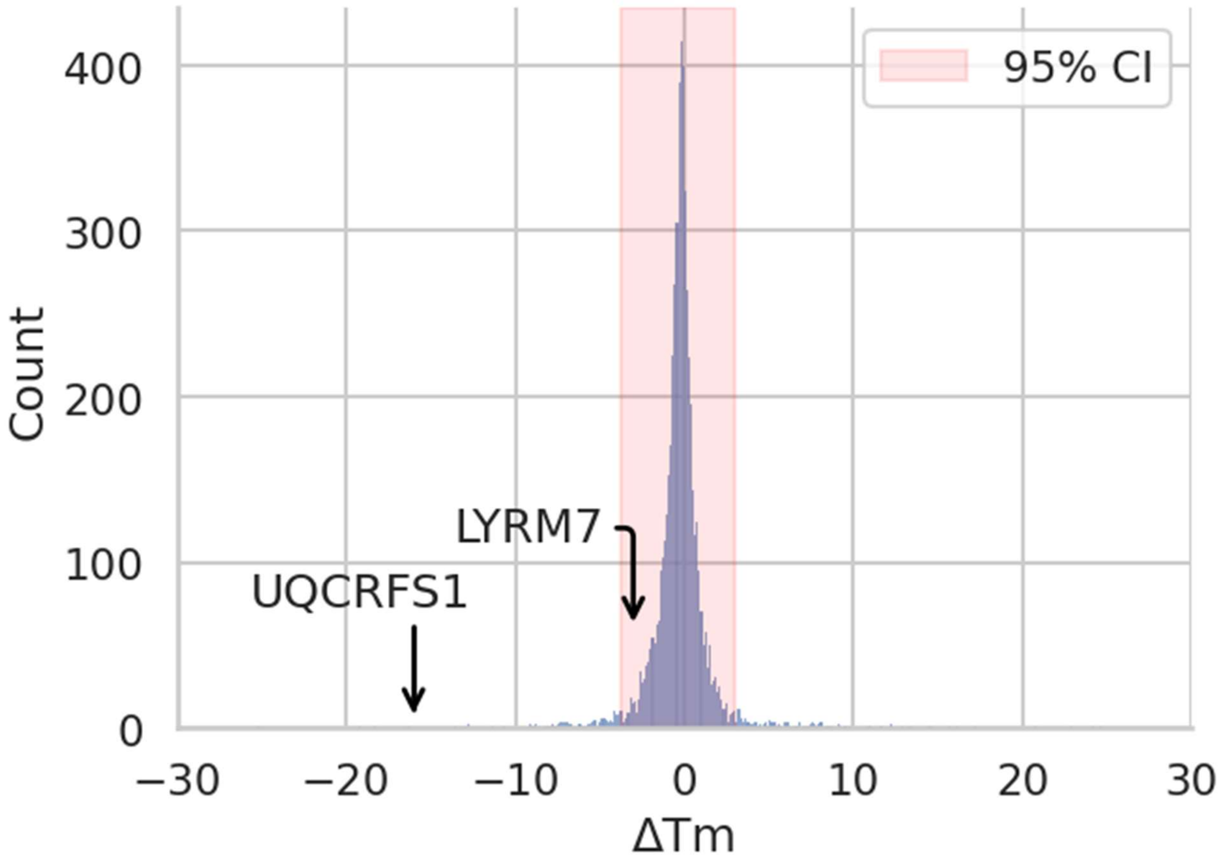

**Supplementary Figure 2.**

**A.**

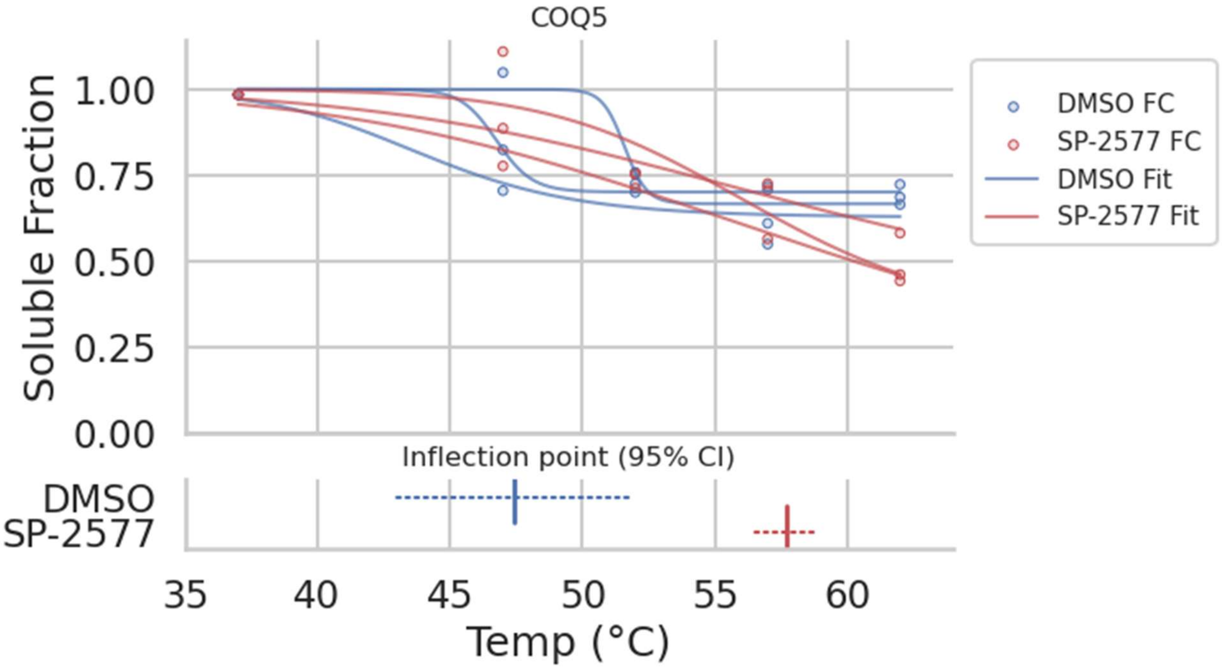


**B.**

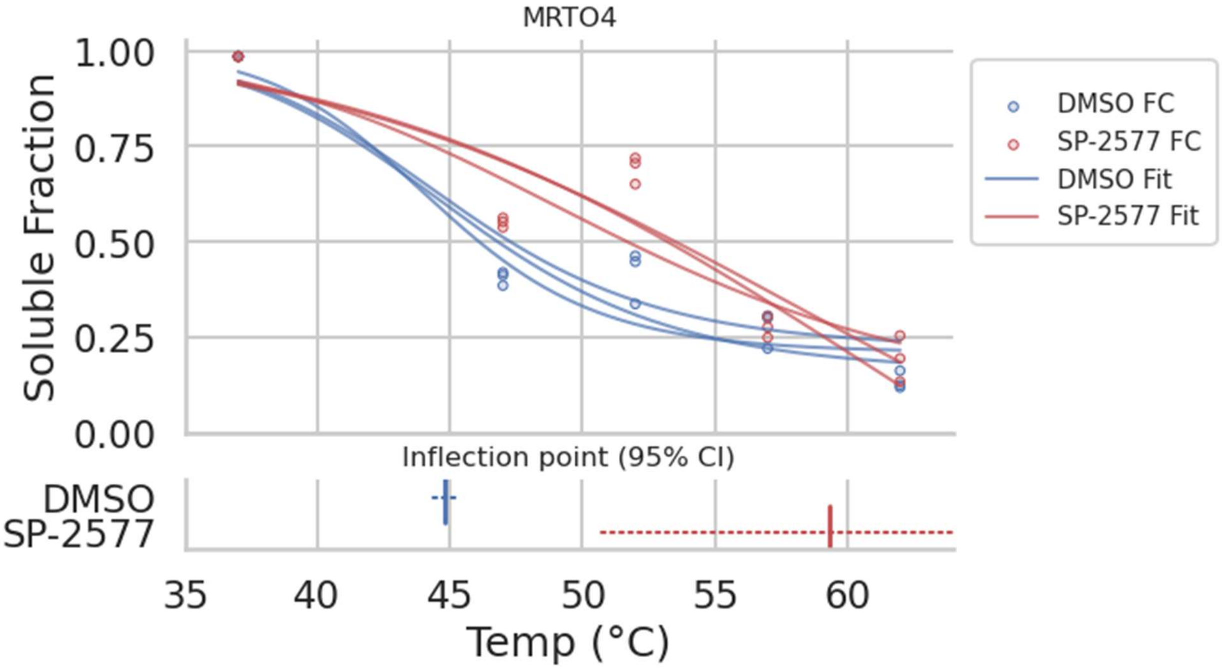

C.

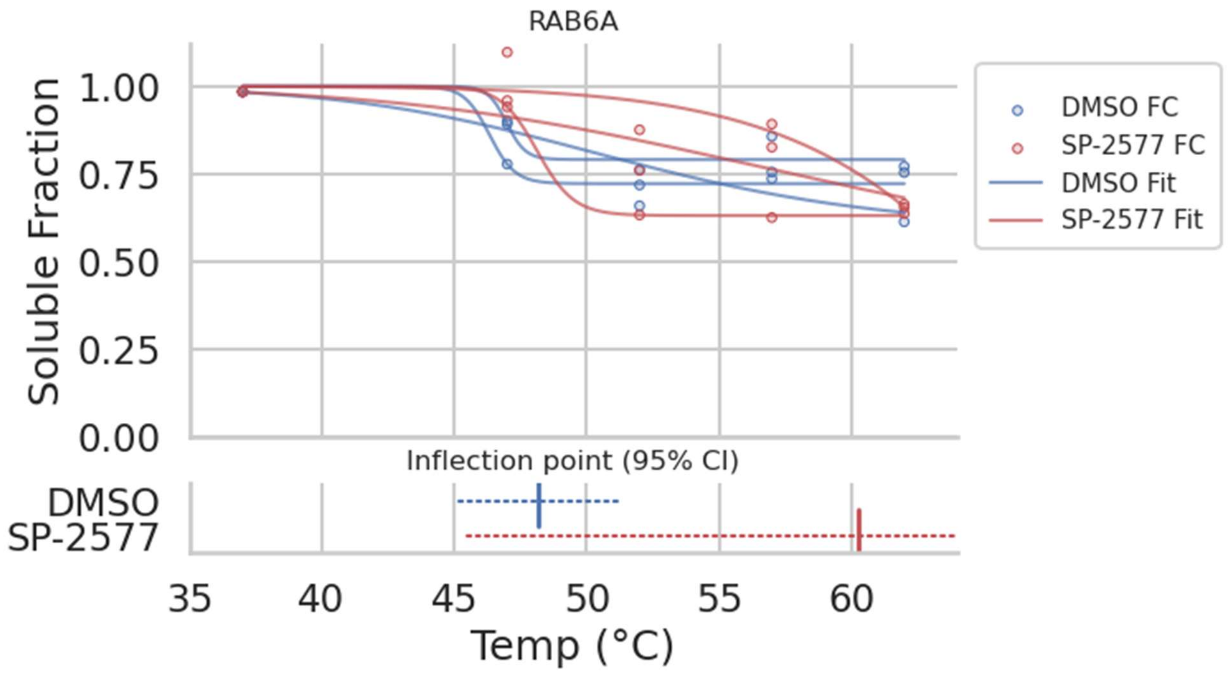

RAB6A
Soluble Fraction
1.00
0.75
0.50
0.25
0.00
DMSO FC
SP-2577 FC
DMSO Fit
SP-2577 Fit
Inflection point (95% CI)
DMSO
SP-2577
35
40
45
50
55
60
Temp (°C)


D.

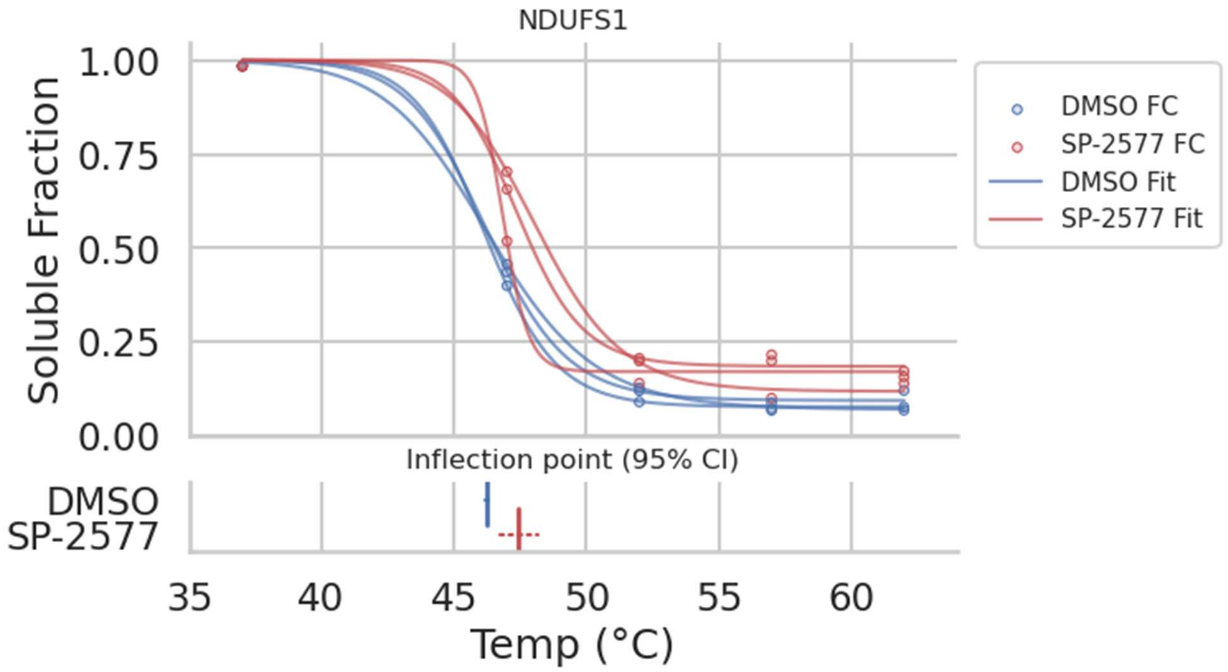

NDUFS1
Soluble Fraction
1.00
0.75
0.50
0.25
0.00
DMSO FC
SP-2577 FC
DMSO Fit
SP-2577 Fit
Inflection point (95% CI)
DMSO
SP-2577
35
40
45
50
55
60
Temp (°C)

**Supplementary Figure 3.**

| °C | 37 | | 38.3 | | 41.4 | | 45.8 | | 51.4 | | 56 | | 58.6 | |
|---|---|---|---|---|---|---|---|---|---|---|---|---|---|---|
| SP-2509 | - | + | - | + | - | + | - | + | - | + | - | + | - | + |

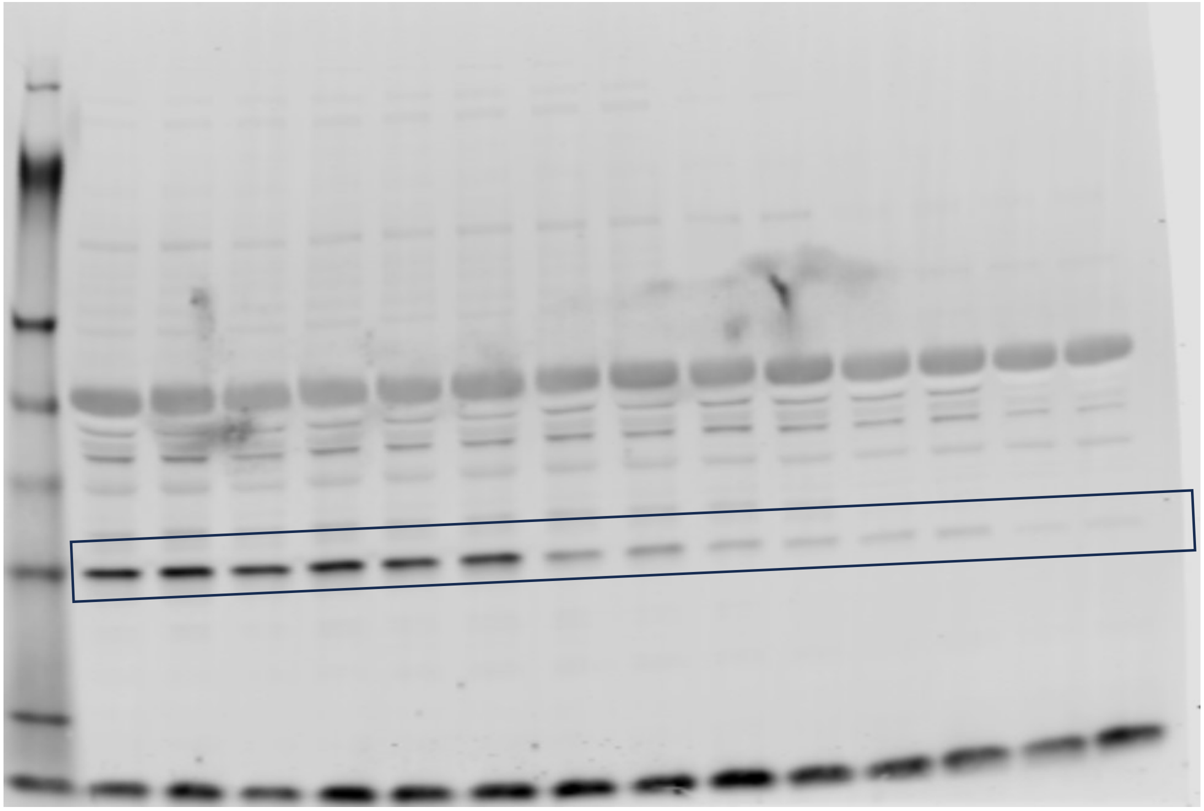

**Supplementary Figure 4.**

| °C | 37 | | 38.3 | | 41.4 | | 45.8 | | 51.4 | | 56 | | 58.6 | |
|---|---|---|---|---|---|---|---|---|---|---|---|---|---|---|
| SP-2509 | **-** | **+** | **-** | **+** | **-** | **+** | **-** | **+** | **-** | **+** | **-** | **+** | **-** | **+** |

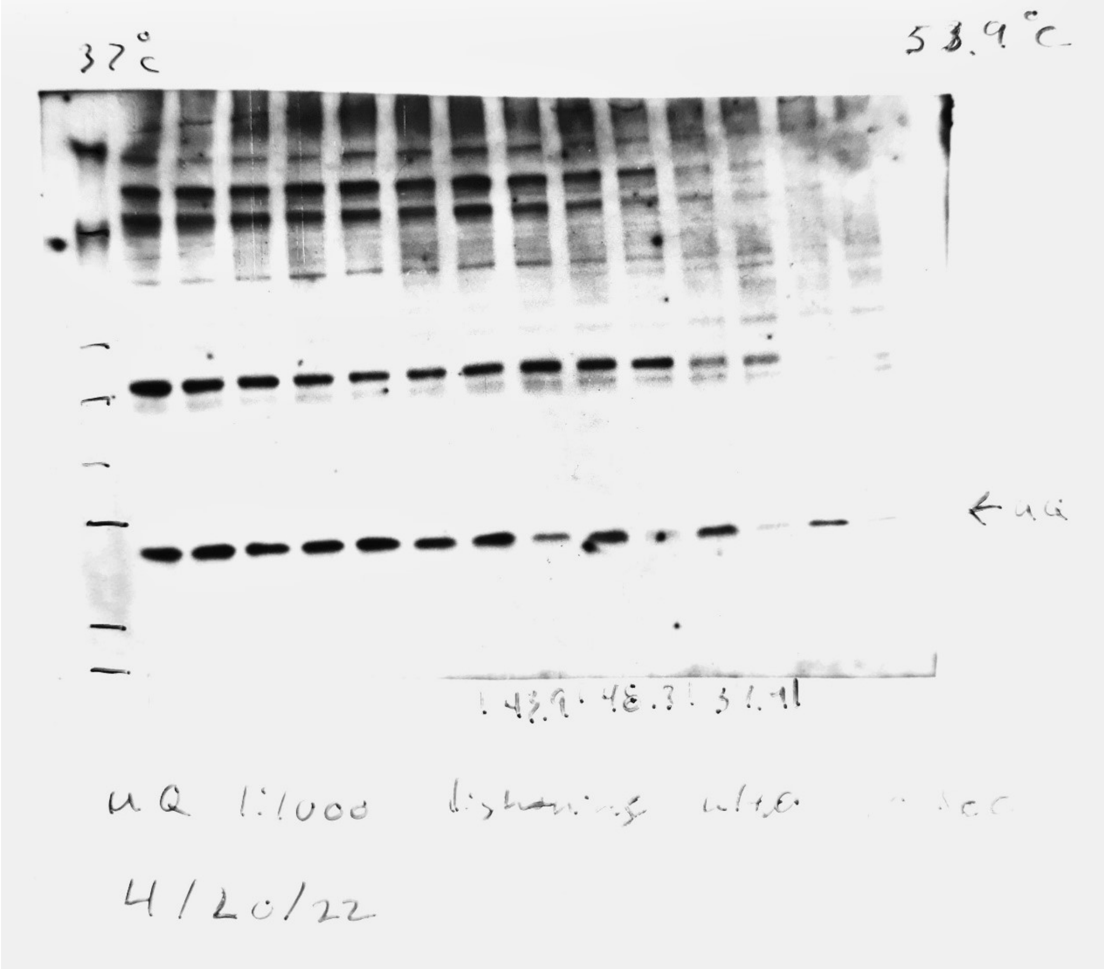


UQCRFS1

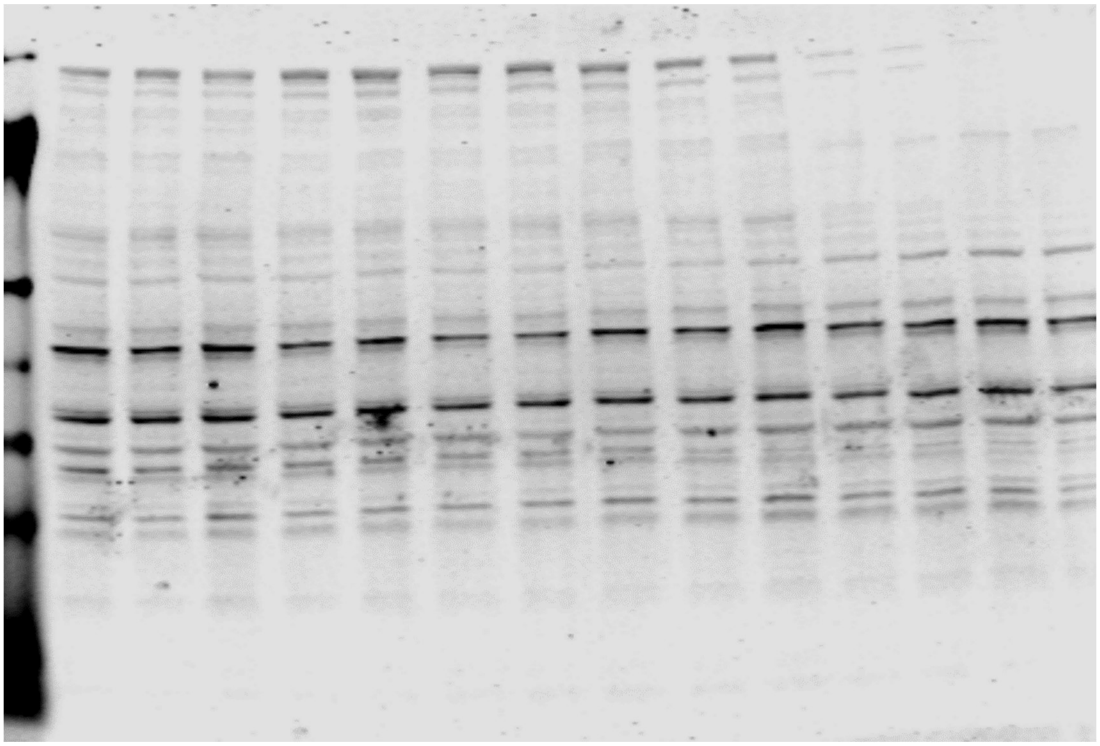

CYC1

**Supplementary Figure 5.**

| °C | 37 | | 38.1 | | 43.9 | | 51.9 | | | 53.9 | | 55 | |
|---|---|---|---|---|---|---|---|---|---|---|---|---|---|
| SP-2509 | - | + | - | + | - | + | - | + | | - | + | - | + |

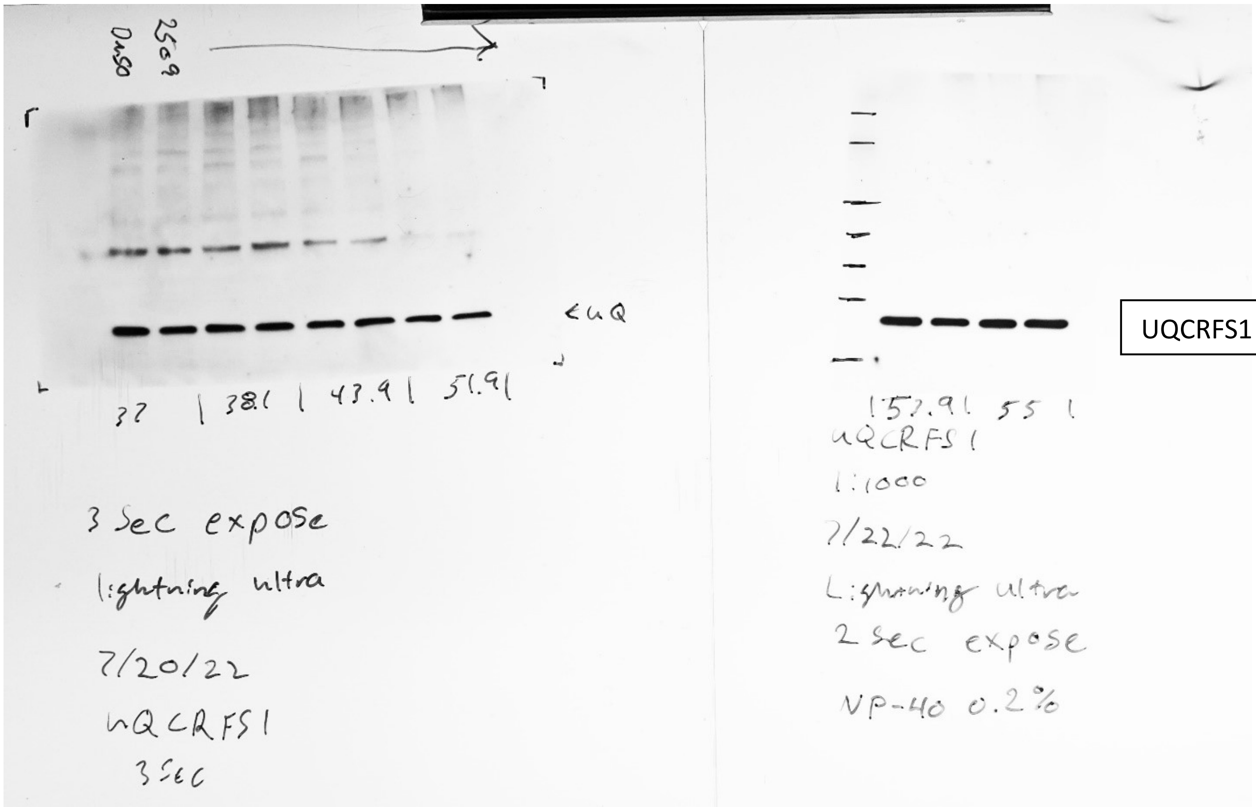


UQCRFS1

| °C | 37 | | 38.1 | | 43.9 | | 51.9 | | | 53.9 | | 55 | |
|---|---|---|---|---|---|---|---|---|---|---|---|---|---|
| SP-2509 | - | + | - | + | - | + | - | + | | - | + | - | + |

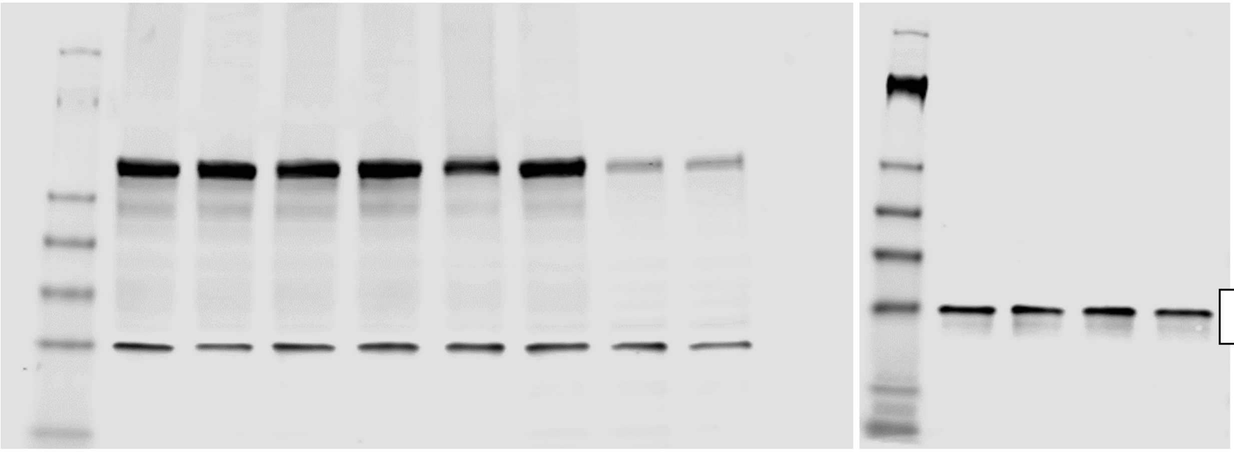

CYC1
